# Biocompatible polymers for scalable production of human neural organoids

**DOI:** 10.1101/2022.03.18.484949

**Authors:** Genta Narazaki, Yuki Miura, Sergey D. Pavlov, Mayuri Vijay Thete, Julien G. Roth, Sungchul Shin, Sarah C. Heilshorn, Sergiu P. Pașca

**Affiliations:** Department of Psychiatry and Behavioral Sciences, Stanford University, Stanford, CA 94305, USA; Institute for Stem Cell Biology and Regenerative Medicine, Stanford University, Stanford, CA 94305, USA; Stanford Brain Organogenesis, Wu Tsai Neuroscience Institute, Stanford, CA 94305, USA; Department of Materials Science and Engineering, Stanford University, Stanford, CA 94305, USA

## Abstract

The generation of neural organoids from human pluripotent stem cells holds great promise in modeling disease and screenings drugs, but current approaches are difficult to scale due to undesired organoid fusion. Here, we develop a scalable neural organoid platform by screening biocompatible polymers that prevent fusion of organoids cultured in suspension. We show that addition of one inexpensive polysaccharide enables straightforward screening of 298 FDA-approved drugs and teratogens for growth defects using over 2,400 cortical organoids.

## Introduction

Human pluripotent stem cell-derived 3D neural culture methods can be used to study human brain development and to investigate neuropsychiatric disease^1–3^. Current approaches to derive neural organoids involve either embedding of individual organoids into extracellular matrices or culturing organoids in suspension^4, 5^. We have previously developed an approach to model human cerebral cortical development using human induced pluripotent stem (hiPS) cell-derived regionalized neural organoids, called human cortical spheroids or human cortical organoids (hCO)^6, 7^. We have shown that hCO, which are derived exclusively in suspension cultures without the use of extracellular matrices, recapitulate features of corticogenesis^8^, can be reliably derived across multiple experiments and lines^9^, and can be used to model environmental and genetic brain disorders^10–14^. Although this suspended culture approach is relatively straightforward to implement, maintaining neural organoids is time-intensive and requires efforts to manually separate and avoid undesired fusion of organoids over time. This spontaneous fusion can often limit the derivation of organoids of uniform size and the scale of experiments.

## Results

To address this issue, we screened biocompatible polymers that can be added to culture medium to prevent fusion and enable culture of multiple organoids in a single well (**Fig. 1a**). We selected 23 candidate polymers based on molecular weight, chemical structure, electrostatic charge, and biocompatibility (see details in **Supplementary Table 1**) and supplemented the neural differentiation medium with each of them starting at day 6. We evaluated the effect of each polymer by culturing five hCO in each well of a 24-well plate for 25 days and then counting the number of remaining hCO in each well (**Fig. 1a**). We found that culture medium supplemented with xanthan gum (XG)-a biocompatible exopolysaccharide widely used in food and pharmaceutical formulations^15^, significantly reduced spontaneous fusion of hCO compared to untreated culture medium (\**P* = 0.01 for untreated hCO versus hCO + XG, \**P* = 0.03 for untreated hCO versus carboxymethyl cellulose 700 in **Fig. 1b**, **Fig.1c** and \*\*\*\**P* = 0.001 for **Fig. 1d**), and yielded hCO of uniform size (**Fig. 1e, f**). Supplementation with XG also prevented hCO fusion with higher efficiency than the commercially available STEMdiff™ Neural Organoid Basal Medium 2 (NOBM) which also prevents organoids fusion (\*\*\*\**P* < 0.0001 for untreated hCO versus hCO + XG, \*\**P* = 0.002 for hCO + XG versus hCO + STEMdiff™ NOBM in **Supplementary Fig. 1a,b**). The cross-sectional area of hCO derived in the presence of XG was smaller than in the untreated condition, which is likely related to the lack of fusion (*P* = 0.99 for day 3, \*\*\*\**p* < 0.0001 for day 10, \*\*\*\**p* < 0.0001 for day 15 for **Supplementary Fig. 1c**). We found that XG-treated hCO displayed ventricular zone-like structures containing SOX2^+^ cells (**Fig. 1g**), and the numbers of neural progenitor (SOX2^+^) cells (**Fig. 1h**, and *P* = 0.72 for **Fig. 1i**) and dorsal forebrain progenitor (PAX6^+^) cells (**Fig. 1j**, and *P* = 0.66 for **Fig. 1k**) were comparable to the untreated hCO condition. XG did not induce cytotoxicity as measured by an LDH cytotoxicity assay (*P* = 0.48 for **Supplementary Fig. 1d**), nor did it induce expression of the apoptotic cell death marker cleaved-caspase 3 (**Supplementary Fig. 1e**). Additionally, hCO cultured with XG did not display significant differences in the expression of the forebrain marker *FOXG1 (P* = 0.53 in **Fig.1l**) or the dorsal cortical marker *EMX1 (P* = 0.20 in **Fig.1l**), and lacked expression of the midbrain marker *EN1 (P* = 0.80 in **Fig.1l**), the medial ganglionic eminence marker *NKX2-1* (*P* = 0.45 in **Fig.1l**), the floor plate marker *FOXA2* (*P* = 0.15 in **Fig.1l**), the hypothalamic marker *RAX* (*P* = 0.39 in **Fig.1l**), or the hindbrain/spinal cord marker *HOXB4* (*P* = 0.45 in **Fig.1l**). Taken together, these data indicate that patterning and differentiation of hCO are not adversely affected by the presence of XG in the culture medium.

**Figure 1.**
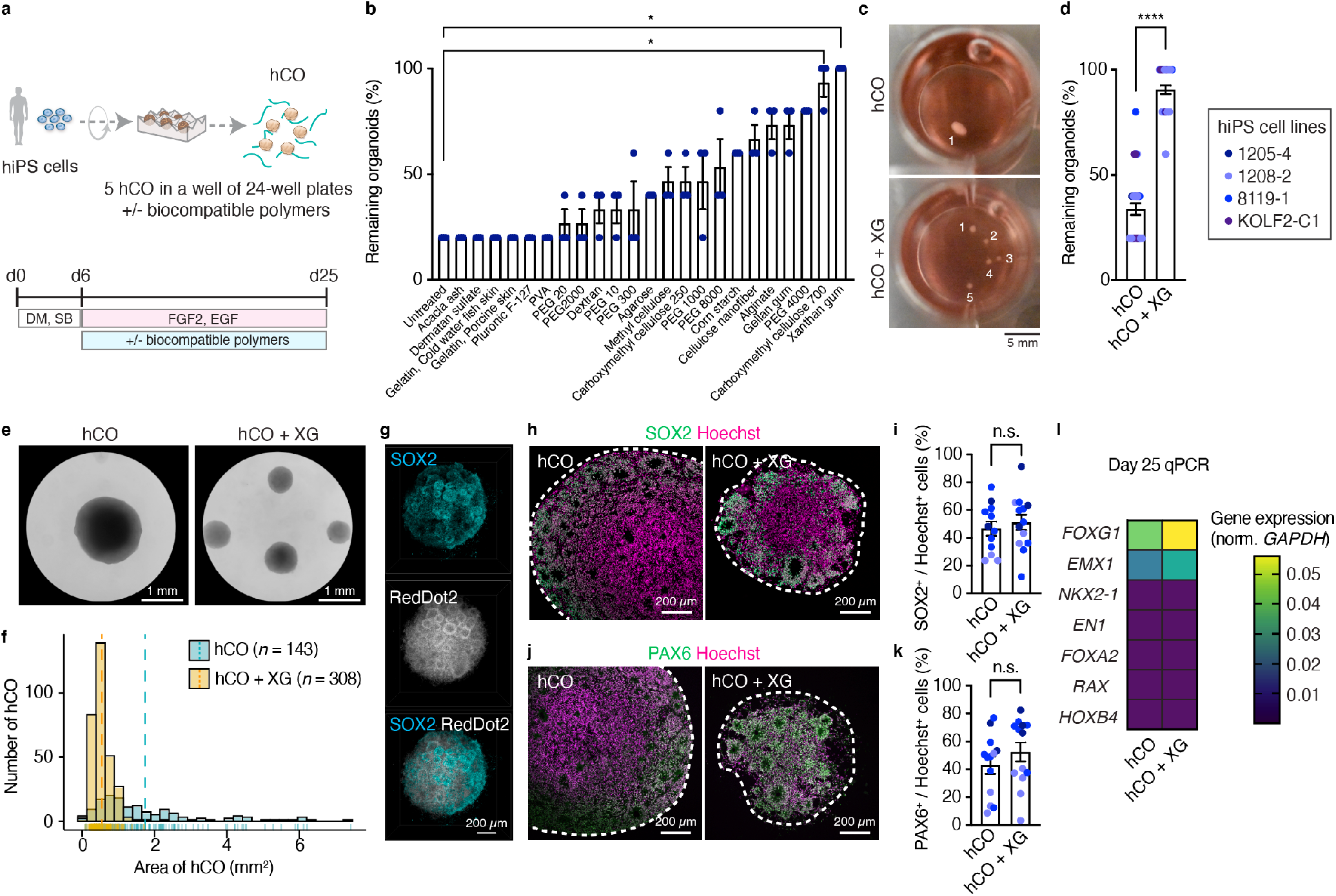
Screening of biocompatible polymers for scalable cortical organoid culture. (**a**) Schematic illustrating the screening process of polymers; 5 hCO were cultured for 25 days in a well of 24 well plates with or without functional polymers, and the number of remaining hCO at day 25 were counted. (**b**) Quantification of remaining organoids at day 25 hCO; *n* = 3 wells for each condition from 1 screening experiment with 1 hiPS cell line. Kruskal–Wallis test, ANOVA result \*\*\*\**P* < 0.0001, \**P* = 0.01 with multiple comparisons for untreated versus xanthan gum, \**P* = 0.03 for untreated versus carboxymethyl cellulose 700. Data show mean ± s.e.m. (**c**) Representative images of untreated hCO (top) and medium with xanthan gum (bottom). Scale bar: 5 mm. (**d**) Quantitative graph of remaining organoids; *n* = 36 wells tested for untreated hCO, *n* = 36 wells tested for hCO + XG from 3 differentiation experiments of 4 hiPS cell lines. Two-tailed unpaired t-test, \*\*\*\**P* < 0.0001. Data show mean ± s.e.m. (**e**) Representative magnified images of untreated hCO and hCO + XG at day 15 of differentiation. Scale bar: 1 mm. (**f**) Histogram showing distribution of organoid area at day 15, blue: untreated hCO, yellow: hCO + XG, dotted lines indicate average for each condition, *n* = 143 organoids for untreated hCO, *n* = 308 organoids for hCO + XG from 4 differentiation experiments including 4 hiPS cell lines. (**g**) 3D immunostaining of CUBIC-cleared hCO cultured with xanthan gum medium. Scale bar: 200 μm. (**h**) Immunostaining for SOX2 (green) and Hoechst (magenta), and (**i**) quantification of SOX2+ / Hoechst+ cells in untreated hCO and hCO + XG at day 25. *n* = 12 organoids for untreated hCO, *n* = 13 organoids for hCO + XG from 2 differentiation experiments of 3 hiPS cell lines. Two-tailed unpaired *t*-test, *P* = 0.72. Scale bar: 200 μm. Data show mean ± s.e.m. (**j**) Immunostaining for PAX6 (green) and Hoechst (magenta), and (**k**) quantification of PAX6^+^ / Hoechst^+^ cells in untreated hCO and hCO + XG at day 25. *n* = 12 organoids for untreated hCO, *n* = 13 organoids for hCO + XG from 2 differentiation experiments of 3 hiPS cell lines. Two-tailed unpaired *t*-test, *P* = 0.66. Scale bar: 200 μm. Data show mean ± s.e.m. (**l**) qPCR of untreated hCO and hCO + XG at day 25; *n* = 7 organoids for untreated hCO, *n* = 7 organoids for hCO + XG from 3 differentiation experiments including 3 hiPS cell lines. Two-tailed Mann-Whitney test, *P* = 0.53 for *FOXG1, P* = 0.80 for *EN1, P* = 0.20 for *EMX1, P* = 0.45 for *NKX2.1, P* = 0.15 for *FOXA2, P* = 0.39 for *RAX, P* = 0.45 for *HOXB4*. Data show mean.

We next verified if XG enables large scale culture and screening of hCO. We compiled a list of 298 drugs that includes FDA-approved compounds for neuropsychiatric conditions and a group of teratogens^16^. We then cultured multiple hCO per well in several 24-well plates and tested the effect of 1 μM of each of the 298 FDA-approved drugs (**Supplementary Table 3**), with DMSO treatment alone as a control, from day 17 to 25 of differentiation (**Fig. 2a**). Importantly, due to the anti-fusion effect of XG, one researcher was able to differentiate and maintain over 2,000 organoids at the same time. To assess morphological changes of hCO, we imaged and analyzed the area of hCO at 8 days after drug exposure and found that several compounds affect the size of hCO (**Fig. 2b, c** and **Supplementary Table 4**). In particular, doxorubicin (Doxo)-a breast cancer drug with known teratogen properties^16, 17^, significantly reduced the size of hCO at 6 days post-exposure (\**P* = 0.02 for day 4, \*\*\*\**P* < 0.0001 for day 6 and 8 for **Fig. 2d**). CUBIC clearing combined with 3D staining of hCO (**Fig. 2e**) confirmed a lower number of SOX2^+^ cells in hCO treated with Doxo (**Fig. 2f** and \*\*\*\**p* < 0.0001 for **Fig. 2g**). We verified this effect using a different batch of Doxo and in multiple hiPS cell lines, and again found growth impairment in Doxo treated hCO (\**P* = 0.02 for **Fig. 2h**), as well as increased cytotoxicity in an LDH cytotoxicity assay (\*\*\**P* = 0.0006 for **Fig. 2i**). Furthermore, we found increased expression of the apoptotic cell marker cleaved caspase-3 (\*\**P* = 0.0064 for **Fig. 2j**), which suggests that Doxo may cause cell death of neural progenitor cells.

**Figure 2.**
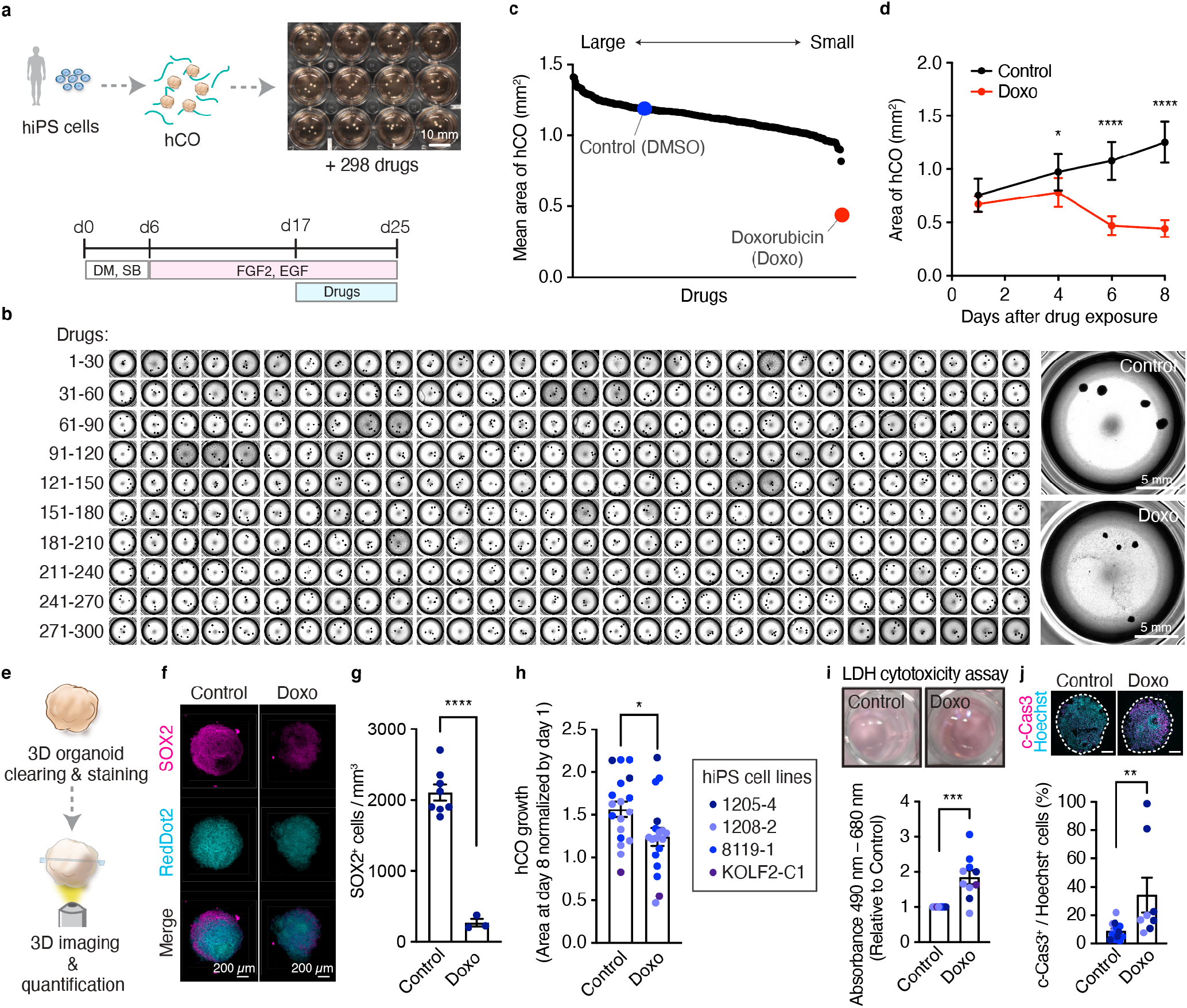
Effects of 298 FDA-approved drugs and teratogens on cortical organoids. (**a**) Schematic showing the screening of 298 drugs in hCO. (**b**) Images of hCO treated with 298 drugs for 8 days. Right top: control, Right bottom: Doxorubicin (Doxo). Scale bar: 5 mm. (**c**) Quantification of organoid size treated with drugs. Data show mean. (**d**) Area of hCO treated with Doxo for 8 days or control; *n* = 8 organoids for control hCO, *n* = *9* organoids for day 1, 4, and *n* = 8 organoids for day 6, 8 of Doxo treated hCO from 1 screening experiment of 1 hiPS cell line. Two-way ANOVA \*\*\*\**P* < 0.0001, \**P* = 0.02 for day 4, \*\*\*\**P* < 0.0001 for day 6 and 8 with Šidák multiple comparison test. Data show mean ± s.d. (**e**) Schematic illustrating 3D clearing and staining of Doxo treated hCO and control. (**f**) 3D immunostaining of CUBIC-cleared hCO cultured with Doxo or control. Scale bar: 200 μm. (**g**) Quantification of SOX2+ cells in 3D-cleared hCO treated with Doxo or control; *n* = 8 organoids for control hCO; *n* = 3 organoids for Doxo treated hCO from 1 screening experiment of 1 hiPS cell line. Two-tailed unpaired *t*-test, \*\*\*\**P* < 0.0001. Data show mean ± s.e.m. (**h**) Organoid growth ratio of hCO treated with Doxo for 8 days or control; *n* = 18 organoids for control hCO, *n* = 18 organoids for Doxo treated hCO from 4 differentiation experiments including 4 hiPS cell lines. Two-tailed unpaired *t*-test, \**P* = 0.02. Data show mean ± s.e.m. (**i**) LDH cytotoxicity assay on day 25 of hCO treated with 1 μM of Doxo for 8 days. Left top image: Control, Right top image: Doxo; *n* = 10 wells for control hCO, *n* = 10 wells for Doxo treated hCO from 3 differentiation experiments including 4 hiPS cell lines. Two-tailed Mann-Whitney test, \*\*\**P* = 0.0006. Data show mean ± s.e.m. (**j**) Immunostaining for cleaved-Caspase-3 (c-Cas3) (magenta) and Hoechst (cyan) in hCO treated with Doxo for 8 days or control at day 25, and quantification of c-Cas3^+^/Hoechst^+^ cells; *n* = 14 organoids for control hCO, *n* = 8 organoids for Doxo treated hCO from 2 differentiation experiments including 3 hiPS cell lines. Scale bar: 200 μm. Two-tailed Mann-Whitney test, \*\**P* = 0.0064. Data show mean ± s.e.m.

## Discussion

In this study, we are describing a key improvement to a neural organoid culture approach we have previously developed that now facilitates scalable differentiation. This is achieved by the simple application of an inexpensive, biocompatible polymer that reduces spontaneous fusions of organoids and thereby enabled one single researcher to differentiate and maintain thousands of organoids in parallel. As a proof of concept, we leveraged this platform to examine the effect of several hundred drugs on neural organoid growth and identified a previously known teratogen compound as potentially being cytotoxic to neural cells. The approach we describe here can be combined with automated culture using a robotic liquid handler or robotic arms to further increase scalability. In principle, this polymer can be used for generating other regionalized neural and non-neural organoids. Taken together, we envision that this simple differentiation approach could substantially increase the applications of organoids for large-scale disease modeling and drug screening.

## Acknowledgements

We thank members of the Pasca laboratory at Stanford University for scientific inputs. This work was supported by the Stanford Brain Organogenesis Big Idea Grant from the Wu Tsai Neurosciences Institute (to S.P.P. and S.C.H.), US National Institutes of Health (NIH) BRAINS Award (MH107800) (to S.P.P.) and R01 EB027171 (to S.C.H.), the NYSCF Robertson Stem Cell Investigator Award (to S.P.P.), the Kwan Research Fund (to S.P.P.), the Coates Foundation (to S.P.P.), the Senkut Research Funds (to S.P.P.), The Ludwig Foundation (to S.P.P.), the Chan Zuckerberg Initiative Ben Barres Investigator Award (to S.P.P.), Stanford Medicine Dean’s Fellowship (to Y.M.), the US National Science Foundation (NSF) awards CBET 2033302 and DMR 2103812 (to S.C.H.), and Stanford Maternal & Child Health Research Institute (MCHRI) Postdoctoral Fellowship (to Y.M.), and the Stanford Bio-X Undergraduate Summer Research Program (to S.D.P.). This paper was typeset with the bioRxiv word template by @Chrelli: www.github.com/chrelli/bioRxiv-word-template

## Author contributions

G.N., Y.M. and S.P.P. conceived the project and designed experiments. G.N. performed the screening experiments for biocompatible polymers and drugs. Y.M. carried out differentiation experiments, 3D clearing and staining, immunocytochemistry and data analyses. S.D.P. conducted 3D clearing, staining and analyzed the images. M.V.T. performed the differentiation experiments, RNA extractions and qPCRs, LDH assay and characterization of organoids. J.G.R., S.S. and S.C.H. selected and prepared biocompatible polymers. Y.M. and S.P.P. wrote the manuscript with input from all authors.

## Materials and Methods

### Characterization and maintenance of hiPS cells

Human induced pluripotent stem (hiPS) cell lines were validated using standardized methods as described previously^9, 18^. Cultures were tested for and maintained Mycoplasma free. A total of 4 control hiPS cell lines were used. KOLF2-C1 hiPS cell line was generated by the Wellcome Trust Sanger Institute. Approval for this study was obtained from the Stanford Institutional Review Board (IRB) panel and informed consent was obtained from all subjects. For maintenance of hiPS cells, the cells were cultured on vitronectin-coated plates (5 μg/mL, Thermo Fisher Scientific, A14700) in Essential 8 medium (Thermo Fisher Scientific, A1517001). Cells were passaged every 4 or 5 days with UltraPure™ 0.5 mM EDTA, pH 8.0 (Thermo Fisher Scientific, 15575020).

### Generation of hCO from hiPS cells

For the generation of human cortical organoids (hCO), hiPS cells were incubated with Accutase^®^ (Innovative Cell Technologies, AT104) at 37°C for 7-10 min and dissociated into single cells. Optionally, 1-2 day before spheroid formation, hiPS cells can be exposed to 1% dimethylsulfoxide (DMSO) (MilliporeSigma, D2650) in Essential 8 medium. To obtain uniformly sized spheroids, AggreWell-800 (STEMCELL Technologies, 34815) containing 300 microwells was used. Approximately 2 x 10^6^ single cells were added per AggreWell-800 well in Essential 8 medium supplemented with the ROCK inhibitor Y-27632 (10 μM, Selleckchem, S1049), centrifuged at 100 g for 3 min to capture the cells in the microwells, and incubated at 37°C with 5% CO_2_. After 24h, organoids consisting of approximately 6,666 cells were collected from each microwell by pipetting medium in the well up and down with a cut P1000 pipet tip and transferred into ultra-low attachment plastic dishes (Corning, 3262) in Essential 6 medium (Thermo Fisher Scientific, A1516401) supplemented with two SMAD pathway inhibitors - dorsomorphin (2.5 μM, Sigma-Aldrich, P5499) and SB-431542 (10 μM, R&D Systems, 1614). On day 6 in suspension the organoids were transferred to neural medium containing Neurobasal™-A Medium (Thermo Fisher Scientific, 10888022), B-27™ Supplement, minus vitamin A (Thermo Fisher Scientific, 12587010), GlutaMAX™ Supplement (1:100, Thermo Fisher Scientific, 35050079), Penicillin-Streptomycin (1:100, Thermo Fisher Scientific, 15070063), and supplemented with 20 ng/mL EGF (R&D Systems, 236-EG) and 20 ng/mL FGF2 (R&D Systems, cat. no. 233-FB). From day 15 of differentiation, media was changed every other day. From day 25, to promote differentiation of the neural progenitors into neurons, the neural medium was supplemented with brain-derived neurotrophic factor (BDNF; 20ng/mL, PeproTech, 450-02), NT3 (20 ng/mL, PeproTech, 450-03). From day 45, only neural medium containing B-27™ Supplement, minus vitamin A (Thermo Fisher Scientific, 12587010) was used for medium changes every 4 days.

### Screening of biocompatible polymers to prevent hCO fusion

At day 6 of differentiation, five hCO were transferred into one well of 24 well plate containing culture media with 0.1% (wt/vol) of polymers (see **Supplementary Table 1**). At day 25, the remaining organoids in each well were imaged using REVOLVE microscope (ECHO), and their number were counted. For comparing to hCO cultured in STEMdiff™ Neural Organoid Basal Medium 2 (hCO + STEMdiff™ NOBM, STEMCELL Technologies, 08620), the remaining organoids in each well were imaged at day 12 on a REVOLVE microscope (ECHO) or a BZX-710 (KEYENCE) with 4x lens.

To prepare the XG-supplemented media, the XG powder, which is difficult to dissolve, was mixed with 100% ethanol. More specifically, to make 250 ml of XG-supplemented Neurobasal medium (0.2% XG [wt/vol], which is a) 2x stock solution), 0.5 g of XG powder was mixed with 2 ml of 100% ethanol, and 248 ml of Neurobasal was then immediately added on top and mixed to avoid the formation of XG clumps. To avoid contamination, dedicated equipment was used inside a biosafety cabinet to weight XG and to mix the medium.

### Cryoprotection and immunocytochemistry

hCO were fixed in 4% paraformaldehyde (PFA)/phosphate buffered saline (PBS) overnight at 4°C. Next day, they were washed in PBS and transferred to 30% sucrose/PBS for 2-3 days until the organoids sink in the solution. Subsequently, they were rinsed in optimal cutting temperature (OCT) compound (Tissue-Tek OCT Compound 4583, Sakura Finetek) and 30% sucrose/PBS (1:1), and embedded. For immunofluorescence staining, 10 or 20 μm-thick sections were cut using a Leica Cryostat (Leica, CM1860). Cryosections were washed with PBS to remove excess OCT on the sections and blocked in 10% Normal Donkey Serum (NDS, Abcam, ab7475), 0.3% Triton X-100 (Millipore Sigma, T9284-100ML), 0.1% BSA diluted in PBS for 1 h at room temperature. The sections were then incubated overnight at 4 °C with primary antibodies diluted in PBS containing 2% NDS, 0.1% Triton X-100. PBS was used to wash the primary antibodies and the cryosections were incubated with secondary antibodies in PBS containing 2% NDS, 0.1% Triton X-100 for 1 h. The following primary antibodies were used for staining: anti-SOX2 (rabbit, Cell Signaling Technology, #3579, 1:300 dilution), and anti-PAX6 (mouse, DSHB, 1:50 dilution), anti-Cleaved Caspase-3 (Asp175) (rabbit, Cell Signaling Technology, #9661, 1:200 dilution). Alexa Fluor dyes, donkey anti-rabbit IgG (H&L) highly cross-adsorbed secondary antibody, Alexa Fluor 488 (Thermo Fisher Scientific, A-21206) and donkey anti-mouse IgG (H&L) highly cross-adsorbed secondary antibody Alexa Fluor 568 (Thermo Fisher Scientific, A10037, 2110843) were used at 1:1,000 dilution, and nuclei were visualized with Hoechst 33258 (Life Technologies, H3549, 10,000 dilution). Cryosections were mounted for microscopy on glass slides using Aquamount (Polysciences, 18606), and imaged on a Leica TCS SP8 confocal microscope. Images were processed in ImageJ/Fiji (Version 2.3.0/1.53f, NIH) and quantified using Imaris (Oxford Instruments, ver. 9.6).

### Real-time qPCR

mRNA from hCO and hCO + XG at day 25 were isolated using the RNeasy Mini kit (Qiagen, 74106) with DNase I, Amplification Grade (Thermo Fisher Scientific, 18068-015). Template cDNA was prepared by reverse transcription using the SuperScriptTM III First-Strand Synthesis SuperMix for qRT-PCR (Thermo Fisher Scientific, 11752250). qPCR was performed using the SYBR™ Green PCR Master Mix (Thermo Fisher Scientific, 4312704) on a QuantStudio™ 6 Flex Real-Time PCR System (Thermo Fisher Scientific, 4485689). Primers used in this study are listed on **Supplementary Table 2**.

### Testing effects of drugs on hCO

At day 17 of differentiation, multiple hCO were transferred into one well of 24-well plate containing neurobasal media with Xanthan gum. These hCO were treated with 1 μM of FDA-approved drugs or teratogens dissolved in DMSO or water (see **Supplementary Table 3**) or only DMSO used as a control. At 1, 4, 6 and 8 days after drug treatment, all hCO in individual wells were imaged using BZX-710 (KEYENCE) with 4x lens, and the area of each organoid was quantified using imageJ/Fiji (Version 2.3.0/1.53f, NIH).

### LDH cytotoxicity assay

At day 25 of hCO culture, 50 μL of culture media from individual wells of culture well plates were collected and the LDH cytotoxicity assay was performed using CyQUANT™ LDH Cytotoxicity Assay (Thermo Fisher Scientific, C20300) following the company’s instruction.

### Clearing and 3D staining of hCO

To optically clear and image hCO, we applied the hydrophilic chemical cocktail-based CUBIC protocol^19^. hCO at day 25 were fixed with a 4% PFA/PBS solution at 4°C overnight. The next day, samples were washed twice with PBS and incubated in Tissue-Clearing Reagent CUBIC-L (TCI, T3740) at 37°C for 2 d. Samples were washed three times with PBS, then nuclei were stained with RedDot2 (Biotium, #40061, 1:150 dilution) in PBS containing 500 mM NaCl at 37°C for overnight. Samples were washed twice with PBS, and once with solution containing 10 mM HEPES, 10% TritonX-100, 200 mM NaCl and 0.5% BSA (HEPES-TSB) at 37°C for 2 hours, and then stained with anti-SOX2 (rabbit, Cell Signaling Technology, #3579, 1:100 dilution) antibody in HEPES-TSB solution at 37°C for 2 d. Stained spheres were subsequently washed twice with 10% Triton X-100 in PBS and once with HEPES-TSB solution for 2 h each and then incubated with a donkey anti-rabbit IgG (H&L) highly cross-adsorbed secondary antibody, Alexa Fluor 488 (Thermo Fisher Scientific, A-21206, 1:300 dilution) in HEPES-TSB solution at 37°C for 2 d. Samples were washed twice with 10% Triton X-100 in PBS for 30 min and once with PBS for 1 h. After washing with PBS, samples were incubated with Tissue-Clearing Reagent CUBIC-R+ (TCI, T3741) at room temperature for 2 d for refractive index matching. CUBIC-cleared hCO were then transferred into a well of a Corning 96-well microplate (Corning, 4580) in 150 μL of CUBIC-R+ solution and imaged using a 10x objective on a Leica TCS SP8 confocal microscope.

### Statistics

Data are presented as Mean ± s.e.m., Mean ± s.d. or only Mean value. Raw data were tested for normality of distribution, and statistical analyses were performed using unpaired *t*-test (two-tailed) or one-way ANOVA tests with multiple comparison tests. Sample sizes were estimated empirically. GraphPad Prism Version 9.3.1 was used for statistical analyses.

### Data availability

The data in this study are available on request from the corresponding author.

**Supplementary Figure 1.**
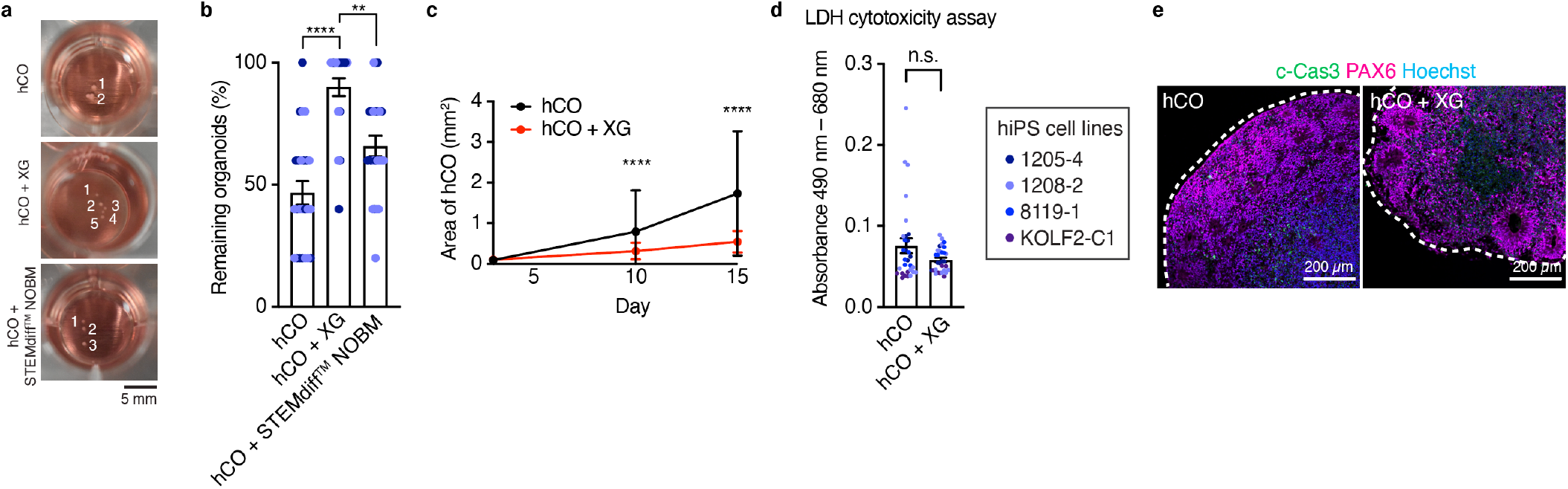
Scalable organoid culture using biocompatible polymers. (**a**) Representative images of untreated hCO (top), xanthan gum-supplemented (hCO + XG, middle) and STEMdiff™ Neural Organoid Basal Medium 2 cultured hCO (hCO + STEMdiff™ NOBM, bottom). Scale bar: 5 mm. (**b**) Graph showing remaining organoids after 5 days of culture; *n* = 24 wells tested for untreated hCO, *n* = 24 wells tested for hCO + XG, *n* = 24 wells tested for hCO + STEMdiff™ NOBM from 2 differentiation experiments of 2 hiPS cell lines. Kruskal–Wallis test, ANOVA result \*\*\*\**P* < 0.0001, \*\*\*\**P* < 0.0001 with multiple comparisons for untreated hCO versus hCO + XG, \*\**P* = 0.002 for hCO + XG versus hCO + ST. Data show mean ± s.e.m. (**c**) Area of hCO at day 3, 10 and 15 cultured with XG or regular culture medium; *n* = 409 organoids for untreated hCO at day 3, *n* = 516 organoids for hCO +XG at day 3, *n* = 208 organoids for untreated hCO at day 10, *n* = 479 organoids for hCO + XG at day 10, *n* = 143 organoids for untreated hCO at day 15, *n* = 308 organoids for hCO + XG at day 15, from 4 differentiation experiments including 4 hiPS cell lines. Two-way ANOVA \*\*\*\**P* < 0.0001, *P* = 0.99 for day 3, \*\*\*\**P* < 0.0001 for day 10, \*\*\*\**P* < 0.0001 for day 15 with Šidák multiple comparison test. Data show mean ± s.d. (**d**) LDH cytotoxicity assay for day 25 hCO or hCO + XG; *n* = 30 wells for untreated hCO, *n* = 30 wells for hCO + XG from 2 differentiation experiments including 4 hiPS cell lines. Two-tailed Mann-Whitney test, *P* = 0.48. Data show mean ± s.e.m. (**e**) Immunostaining of cleaved-caspase 3 (c-Cas3) in day 25 untreated hCO and hCO + XG. Scale bar: 200 μm.

